# The impact of host metapopulation structure on the population genetics of colonizing bacteria

**DOI:** 10.1101/027581

**Authors:** Elina Numminen, Michael Gutmann, Mikhail Shubin, Pekka Marttinen, Guillaume Méric, Willem van Schaik, Teresa M. Coque, Fernando Baquero, Rob J.L. Willems, Samuel K. Sheppard, Edward J. Feil, William P. Hanage, Jukka Corander

**Affiliations:** Department of Mathematics and Statistics, University of Helsinki, Helsinki, Finland; Helsinki Institute for Information Technology HIIT, Department of Information and Computer Science, Aalto University, Finland; College of Medicine, Swansea University, Institute of Life Science, Swansea, UK; Department of Medical Microbiology, University Medical Center Utrecht, Utrecht, The Netherlands; Department of Microbiology, Ramón y Cajal University Hospital, Madrid, Spain; Department of Biology and Biochemistry, University of Bath, Claverton Down, Bath, UK; Center for Communicable Disease Dynamics, Harvard School of Public Health, Boston, Massachusetts, USA

**Keywords:** Bacterial evolution, genetic structure, migration, population dynamics

## Abstract

Many key bacterial pathogens are frequently carried asymptomatically, and the emergence and spread of these opportunistic pathogens can be driven, or mitigated, via demographic changes within the host population. These inter-host transmission dynamics combine with basic evolutionary parameters such as rates of mutation and recombination, population size and selection, to shape the genetic diversity within bacterial populations. Whilst many studies have focused on how molecular processes underpin bacterial population structure, the impact of host migration and the connectivity of the local populations has received far less attention. A stochastic neutral model incorporating heightened local transmission has been previously shown to fit closely with genetic data for several bacterial species. However, this model did not incorporate transmission limiting population stratification, nor the possibility of migration of strains between subpopulations, which we address here by presenting an extended model. The model captures the observed population patterns for the common nosocomial pathogens *Staphylococcus epidermidis* and *Enterococcus faecalis,* while *Staphylococcus aureus* and *Enterococcus faecium* display deviations attributable to adaptation. It is demonstrated analytically and numerically that expected strain relatedness may either increase or decrease as a function of increasing migration rate between subpopulations, being a complex function of the rate at which microepidemics occur in the metapopulation. Moreover, it is shown that in a structured population markedly different rates of evolution may lead to indistinguishable patterns of relatedness among bacterial strains; caution is thus required when drawing evolution inference in these cases.

## Introduction

Bacteria colonizing multicellular hosts are organized in a hierarchy of local interconnected subpopulations forming a complex metapopulation as a whole. The subpopulations can range in scale from discrete intracellular colonies residing within a single host cell to pervasive strains circulating among hosts across cities, countries and continents(Fraser et al., 2009). Although most bacteria are harmless or even advantageous to their host organisms, some cause infectious disease, and understanding the evolutionary dynamics and the factors producing the genetic variation of pathogen populations is important for combatting disease emergence and spread.

Previous work has demonstrated that a simple model of stochastic microepidemics arising from repeated sampling of localized transmission chains, can explain genotypic variation in local surveillance data from several common human pathogens(Fraser et al., 2005; Hanage et al., 2006), under an assumption that all isolates are equally fit (neutrality). In these studies, populations were characterized by a simple measure of the level of genotype relatedness known as the allelic mismatch distribution, where isolates with more shared alleles are considered to be more closely related. These comparisons have been widely used in classical ecology and population genetics and different patterns in the mismatch distribution can be associated with various factors contributing to the population structure, including: population growth(Harpending, 1994; Rogers and Harpending, 1992), selection(Bamshad et al., 2002), and host contact network structure(Plucinski et al., 2011). The mismatch distribution has also been used to detect deviations from neutrality or constant population size(Mousset et al., 2004) and for inference about bacterial recombination rates(Hudson, 1987).

Population structure is one of the most studied phenomena in population genetics, both from the theoretical and applied perspective(Ewens, 2004; Hartl and Clark, 2007). Nevertheless in the case of bacteria limited knowledge exists about the effects of population structure arising from multiple host organisms such as human and different animal species or other, often poorly defined and understood, ecological patches. The main reason for this is simultaneously accounting for the major phenomena known to impact evolution of bacterial pathogen populations, such as recombination, clonal expansion, as well as migration, which for example may be caused by anthroponosis and zoonosis when multiple different host organisms are colonized by the same bacterial species. This hampers both theoretical derivation of limit results for such models and empirical fitting due to likelihood equations not being available in closed form. Fraser et al. solved the likelihood intractability arising from microepidemics by using a stochastic mixture distribution to account for the increase in the probability of sampling identical strains from the same transmission chain (Fraser et al., 2005). An analogous approximation technique has later been independently introduced in a more general ecological setting and it is known as the *synthetic likelihood* (Wood, 2010).

To improve understanding of the evolutionary dynamics of structured bacterial populations, we employ a simulation-based approach to neutral models that can account for the multiple stochastic forces impacting the genetic diversity that persists over time. By capturing both a heterogeneous span of microepidemics and migration events across the boundaries limiting transmission between subpopulations, we characterize the expected behavior of the metapopulations as a whole. This provides an opportunity to explore the limits of inferring the vital model parameters from genetic surveillance data, and gives novel insight into the emergence of important human pathogens.

## Materials and Methods

### Model

We consider an infinite alleles model for a finite haploid population with *N* individuals and discrete generations, where the reproduction takes place by random sampling of *N* individuals from the current generation to the next generation(Ewens, 2004). When the population is assumed structured, the subpopulation sizes are indexed by *N*_1_, *N*_2_. The parameters which may vary across subpopulations are indexed accordingly. Mutations are introduced per generation by a Poisson process with the rate *θ* = *μNτ,* where *μ* is the per locus mutation rate and *τ* is a scaling factor representing the generation time in calendar time. In all subsequent work we set *τ* = 1, unless otherwise mentioned. We assume that each individual is characterized by a genotype comprising alleles at *L* unlinked loci, where a mutation event at any locus always introduces a novel allele. Recombination between randomly chosen genotypes occurs at any locus according to a Poisson process with the rate defined as *ρ* = *rNτ,* where *r* is the rate per locus in relation to the mutation rate. In our simulations we simulated the population until allelic diversity reached equilibrium.

Microepidemics are modeled as doubly stochastic events, with the frequency of new microepidemics per generation following a Poisson distribution with mean *μNτ.* The size of each microepidemic has a Poisson distribution with mean *γ.* Each micropidemic is generated independently similar to the assumptions in Fraser et al. such that first a single individual is randomly chosen, after which its genotype is propagated to *Y* randomly chosen other individuals such that *Y* has Poisson distribution with mean *γ.* When the population is stratified, the microepidemic rates of the subpopulations are denoted by *ω*_1_, *γ*_1_ and *ω*_2_*, γ*_2_ respectively. Migration between subpopulations is a Poisson process with the rates τ*N*_1_*m*_12_, τ*N*_2_*m*_21_ per generation, where the first subindex of the parameters *m*_12_*, m*_21_ defines the source and the second subindex the target subpopulation. In migration events genotypes of a Poisson distributed number of randomly chosen individuals from the source population replace the genotypes of randomly chosen individuals in the target population. In our simulations the events were generated in the following order: reproduction mutation, recombination, microepidemics and migration at each generation. In all the reported results each subpopulation size was *N* = 2000, unless otherwise indicated. Medians and 95% confidence intervals for the allelic mismatch distributions were obtained by recording the population state every 100th generation after initial 500 generations until 20000 generations, and using these values to calculate the corresponding quantiles of the mismatch probabilities.

### Data and processing of genotype networks

eBURST networks for the populations were produced using default settings(Feil et al., 2004). Turner et al. demonstrated that eBURST provides a robust recapitulation of the genetic relatedness of strains in a bacterial population based on the MLST resolution(Turner et al., 2007). To quantify details of the networks we calculated genotype degree distributions and distributions of geodesic distances between pairs of genotypes, which are standard measures of network topology(Goh et al., 2002). MLST isolate data were accessed (September 15, 2014) from the following databases: http://efaecalis.mlst.net/ *(E. faecalis),* http://efaecium.mlst.net/ *(E. faecium),* http://saureus.mlst.net/ *(S. aureus),* and (May 10, 2015) from: http://sepidermidis.mlst.net/ *(S. epidermidis).*

## Results

We extended the microepidemic infinite alleles model with mutation and recombination rates previously proposed by Fraser et al. (Fraser et al., 2005) to incorporate population stratification, whereby genotypes are free to move between subpopulations at a defined rate. In addition, rather than using a single microepidemic parameter to describe localized transmission (Fraser et al., 2005), we introduced two parameters modulating the distributions of both the frequency and sizes of the transmission clusters in stochastic fashion. Our microepidemic infinite alleles migration model (MIAMI) can thereby encompass a wide variety of evolutionary and ecological parameter space. Since the resulting patterns of genetic variation reflect a complex function of several factors, we consider first a model without population stratification to delineate the influence of each of the model components.

The frequency distribution of the number of allelic mismatches between pairs of genotypes is a classical approach to describe the distribution of genetic variation within a population (Fraser et al., 2005). Depending on the interplay of several factors, a population may either have a peaked or flat equilibrium distribution over the space of summary statistics, such as the allelic mismatch distribution (Fig. 1). For lower mutation rates, high *r/m* will lead to bell-shaped mismatch distributions, since recombination acts as a cohesive force keeping genetic variation together as a cloud in the space of possible genotypes(Fraser et al., 2007). The mismatch distribution becomes less sensitive to changes in recombination rate and the equilibrium distribution becomes more peaked when the mutation rate increases (Fig. 1).

**Fig. 1.**
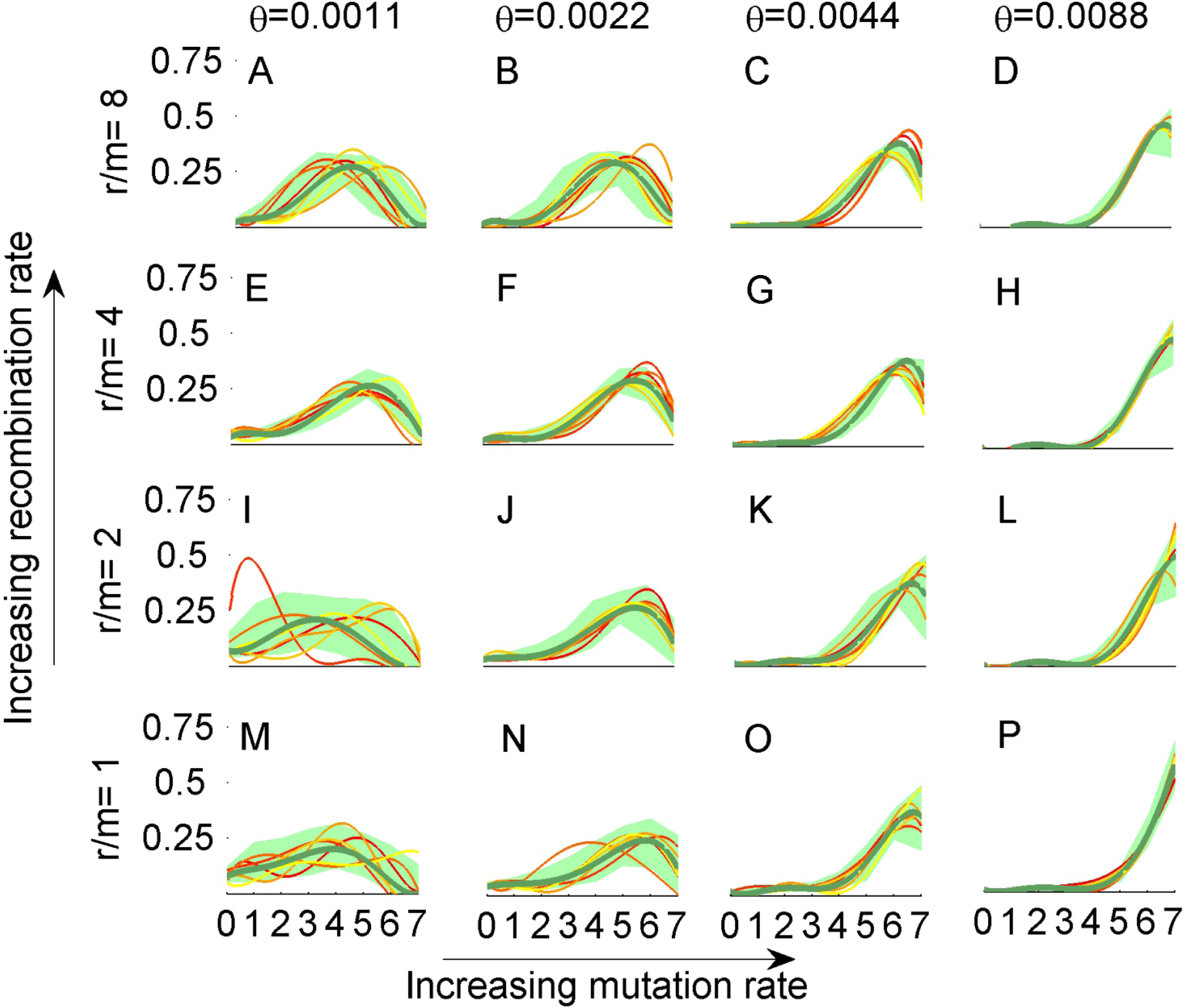
Allelic mismatch distributions for combinations of mutation and recombination rates in a population with *N* = 3000. Bold line in green shows the mean mismatch probability over 20000 generations, sampled at intervals of 100 generations. The green shaded area shows the 95% confidence interval and the colored lines are examples of mismatch distributions at random time points. Vertical axis in each panel shows the probability mass associated with the points of the curves across the values on the horizontal axis. Distributions are shown as continuous curves for visual clarity.

Fig. 2 shows the impact of heightened localized transmission (microepidemics) on genetic relatedness visualized using eBURST (Feil et al., 2004; Francisco et al., 2009) and the allele mismatch distribution. The rate of mutation and homologous recombination varies among bacterial pathgoens and this can have a marked effect on the population structure. To model the interplay of these two important factors at different levels, four evolutionary scenarios were considered: low mutation and recombination rate (A), mutation dominates (B), recombination dominates (C), both mutation and recombination effects are sizeable (D). If mutation dominates over recombination (Fig. 2,B), microepidemics do not lead to as pronounced changes in the relatedness pattern as in the situation where both mutation and recombination rates are low (Fig. 2,A). Interconnected clusters do emerge under a high rate of recombination, often spanning across large parts of the entire population (Fig. 2,C). The variability of the mismatch distribution at the equilibrium becomes elevated under all regimes of baseline parameter values when microepidemics occur at a frequent rate, as illustrated by the broader confidence intervals (Fig. 2,A-D). Both the frequency and size distribution of the individual microepidemics influence how much probability mass is shifted towards identical genotypes, but the change is also influenced by mutation and recombination rate parameters (Supplementary Fig. 1).

**Fig. 2.**
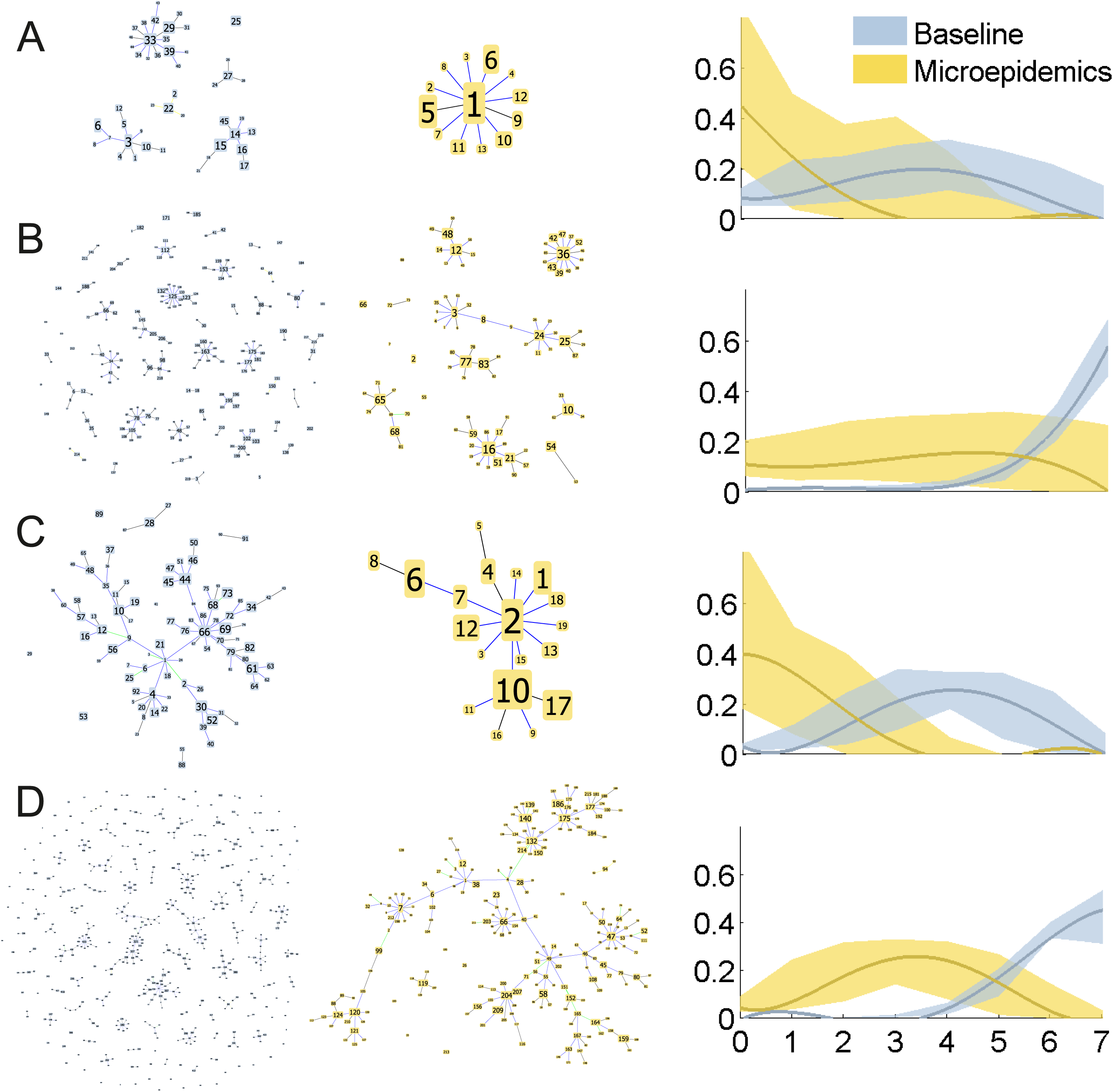
eBURST networks and mismatch distributions for a population without (grey) and with (yellow) microepidemics, where *ω* = 27, *γ* = 16. The 95% confidence intervals are shown by shaded areas and are defined as in Fig. 1. The mutation and recombination parameters used are: 0.0011, 1 (A), 0.0088, 1 (B), 0.0011, 8 (C), 0.0088, 8 (D).

The effect of migration rate on the allelic mismatch distribution within a subpopulation is a complicated function of mutation, recombination and microepidemic rates in a structured population, even if there are only two subpopulations (Fig. 3). We studied the combinations in which a subpopulation undergoes microepidemic expansions at a moderate rate and is coupled with another subpopulation where the rate varies from zero to twice that of the first subpopulation. An increase of the migration rate between the two subpopulations by an order of magnitude leads either to a substantial decrease of the genotypic diversity (Supplementary Fig. 2, i), an increase in the genotypic diversity (Supplementary Fig. 2, a), or to no change at all (Supplementary Fig. 2, e), depending on whether the subpopulation considered as a source experiences more, less, or an equal amount of the microepidemics, compared with the target subpopulation. The effect of migration remains equally complex for the between-subpopulations allelic mismatch distribution, which is insensitive to a change in the migration rate by an order of magnitude for many combinations of subpopulation dynamics (Supplementary Fig. 3). Population stratification combined with asymmetric migration rates can produce patterns of relatedness which are otherwise unlikely under the neutral model (Supplementary Fig. 4). For example, in all our simulations a characteristic U-shaped allelic mismatch distribution only arose when the migration rate was highly asymmetric and one subpopulation experienced considerable microepidemics while the other one had none (Supplementary Figs. 5,6).

**Fig. 3.**
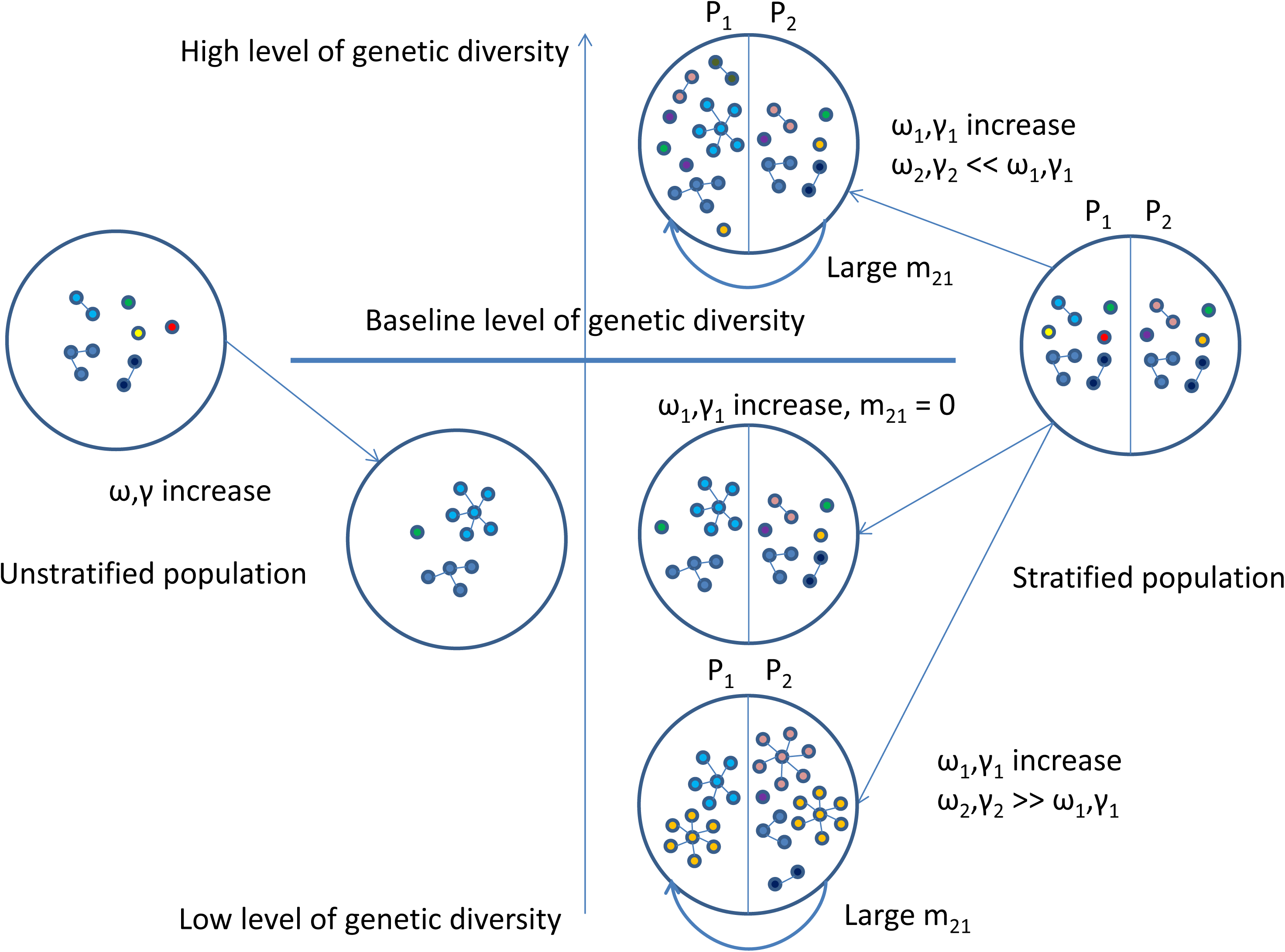
Schematic illustration of the combined effect of microepidemics and migration. The population on the left is unstratified, in which case increasing rate (ω) and size (γ) of microepidemics lead to decreased genetic variation. In a stratified population with two subpopulations (P_1_, P_2_) the effect of increasing microepidemics (ω_1_, γ_1_) on genetic diversity in subpopulation P_1_ depends both on the microepidemics in subpopulation P_2_ (ω_2_, γ_2_) and on the migration rate (m_21_). The case with m_21_ = 0 leads to identical decrease of genetic variation as in an unstratified population. The notation “<<” is used to indicate that the parameters on the left side of the double inequality are much smaller than those on the right side.

**Fig. 4.**
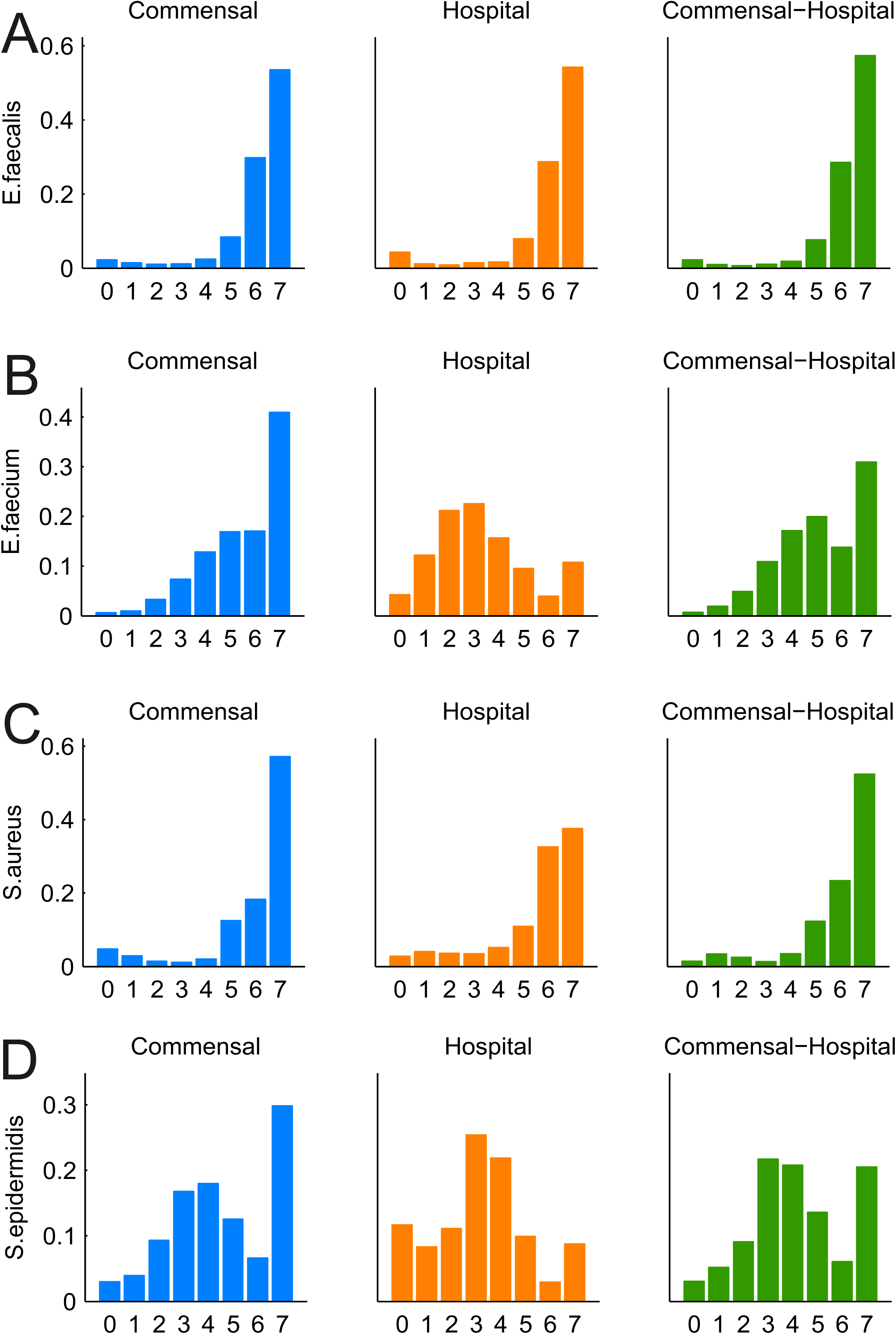
Mismatch distributions of commensal and hospital subpopulations of four common nosocomial bacterial pathogens. The right-most column shows the between-subpopulation mismatch distributions.

To obtain an analytical insight to the joint effect of microepidemic and migration rates on genotypic diversity, we considered how the equilibrium probability of identical genotypes is affected by introducing a change to the subpopulation based on either mechanism. Fraser et al. derived the equilibrium probability of identical genotypes at *L* unlinked loci, under the assumption of no microepidemics(Fraser et al., 2005), which equals 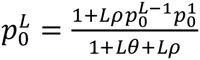 Here *θ* = *2μN*, where *μ* is the per locus mutation rate and *N* is the population size. Furthermore, the recombination rate is defined as *ρ* = 2*rN*, where *r* is the rate per locus in relation to the mutation rate. Since this extension of the classical equilibrium result by Kimura to allow for recombination is based on the assumption that in any generation only a single event occurs, Fraser et al. handled the effect of microepidemics on a population at equilibrium implicitly by introducing a probabilistic mixture where a single parameter represents the increase in the probability 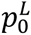 caused by microepidemics. Consistent with this, we quantify the change in the probability of identical strains by evaluating the expectation of the effect of microepidemic and migration events when allowed at the equilibrium of a simpler population experiencing only mutation and recombination events.

Consider first the effect of stochastic microepidemics occurring in a single generation. The expected number of identical genotype pairs arising from them equals (*γ* + 1)^2^*Nω*, where *ω* is the scaled rate at which microepidemics occur per generation and *γ* is the expected size of each microepidemic (Methods). The expected contribution to the probability of homozygous strains is then 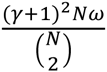 which is an increasing function of both the expected size and rate of microepidemics. Next, consider two subpopulations of sizes *N*_1_*, N*_2_, which at equilibrium become connected with migration rates *N*_1_*m*_12_*, N*_2_*m*_21_, respectively, in addition to the effect of introducing microepidemics (Methods). Each subpopulation is assumed to have its own set of parameters 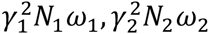 governing the extent of microepidemics. Assume now that the subpopulations are of equal size *N*_1_ = *N*_2_. Then, the expected contribution to the probability of identical strains in subpopulation 1 by an increase in the migration rate *m*_21_ depends on whether 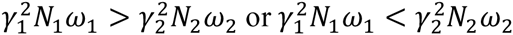, since larger and more frequent microepidemics in subpopulation 2 will increase the probability that the genotypes migrating to subpopulation 1 are identical to each other. Conversely, increased migration from subpopulation 2 will have expected effect of decreasing the probability when the extent of microepidemics in subpopulation 2 is smaller than in subpopulation 1. A difference in the sizes of the subpopulations can further amplify these effects since the rates of events are relative to them.

Global surveillance data based on MLST typing for several common nosocomial bacterial pathogens (*S. aureus, S. epidermidis, E. faecalis, E. faecium*) generally match well with the expected shape of the allelic mismatch distribution for the considered archetypical population types (Fig. 4). eBURST diagrams provide additional insight into the structure of these populations (Fig. 5). *S. aureus* is known to have a very low recombination rate(Everitt et al., 2014) and its population structure is mainly shaped by a combination of mutation rate and intensive clonal expansion of distinct genotypes (Fig. 5, C). Conversely, its sister species *S. epidermidis* displays the bell-shaped mismatch distribution typical for organisms with high recombination rate(Meric et al., 2015) (Fig. 4, D) and a large connected network of related genotypes (Fig. 5, D). The numerous distinct clusters with short distances to the ancestral genotype observed in *S. aureus* population (clonal complexes with single-locus variants) were not accurately predicted by the model, despite of an extensive search over the parameter space. The main deviance arose from the inability to recapitulate a large number of descendant genotypes connected with each single ancestral genotype. The most closely matching neutral model predicts instead invariably that several further branches emerge from these descendants during the timescale at which genotype clusters themselves emerge.

**Fig. 5.**
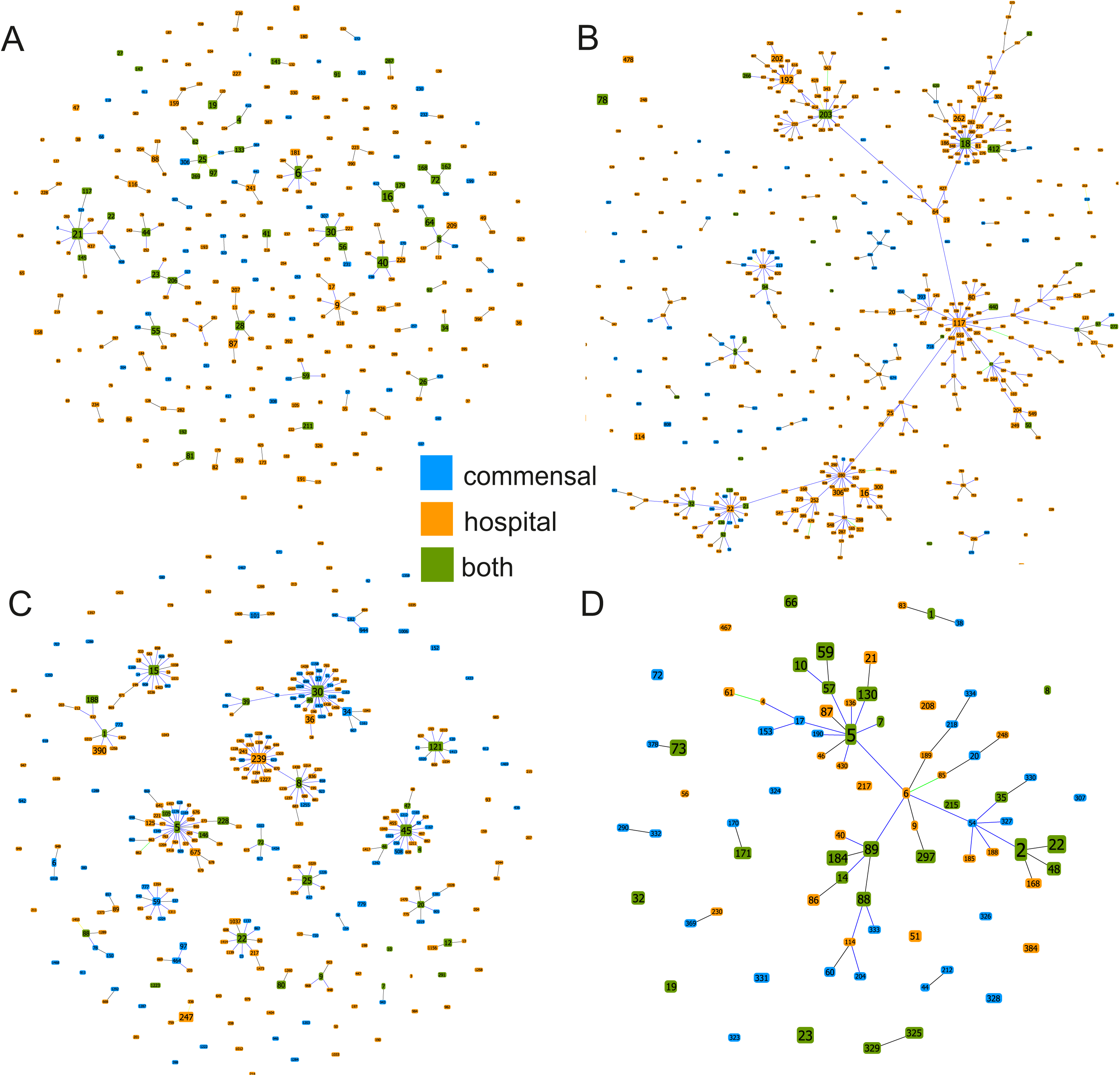
eBURST networks of the isolates used to calculate the mismatch distribution in Fig. 4; *E. faecalis* (A), *E. faecium* (B), *S. aureus* (C), *S. epidermidis* (D).

Contrasting the population structures of *E. faecium* and *E. faecalis* reveals marked differences, where *E. faecium* forms large networks of related genotypes characteristic of highly recombinogenic bacteria (Fig. 5, B) (Turner et al., 2007), despite a relatively low estimated recombination rate(de Been et al., 2013). *E. faecalis* shows only limited clustering of genotypes (Fig. 5, A) and a mismatch distribution typical for a population dominated by mutation, with a slight increase of identical genotype pairs due to localized hospital transmission (Fig. 4, A).

The model parameter configurations leading to matching characteristics between the observed and simulated population structure are given in Table 1 for the two species where the neutral model recapitulates the surveillance data well (*S. epidermidis, E. faecalis*). We compared genotype networks using the standard measures of degree distribution and geodesic distances between nodes and found a considerable agreement between the data and the simulations (Table 1, Supplementary Fig. 7,8).

**Table 1.** Population characteristics of genotype relatedness for real and simulated data.

|  | *S. aureus* | *S. epidermidis* | *E. faecalis* | *E. faecium* |
| --- | --- | --- | --- | --- |
| *N* commensal | 555 | 120 | 225 | 126 |
| *N* hospital | 543 | 264 | 1003 | 1534 |
| Mean degree | 3.14 | 1.95 | 1.12 | 4.03 |
| Max degree | 34 | 10 | 9 | 47 |
| Mean geodesic distance | 2.15 | 3.54 | 1.82 | 4.12 |
| Max geodesic distance | 5 | 7 | 5 | 12 |
| Simulation settings | Not matching | $\omega_1 = 45, \gamma_1 = 30, \omega_2 = 10, \gamma_2 = 10, m_{12} = 0.01, m_{21} = 0.001, \theta = 0.0704, r/m = 2$ | $\omega_1 = 10, \gamma_1 = 20, \omega_2 = 15, \gamma_2 = 20, m_{12} = 0.001, m_{21} = 0.001, \theta = 0.198, r/m = 1$ | Not matching |
| Mean degree |  | 1.67 | 1.08 |  |
| Max degree |  | 20 | 13 |  |
| Mean geodesic distance |  | 3.1 | 1.94 |  |
| Max geodesic distance |  | 8 | 6 |  |

## Discussion

Previously described neutral models specified by mutation and recombination rate in combination with microepidemics show a close fit to observed genotype survey data for several commensal and pathogenic bacteria. This holds true for both short-term population evolution dominated by the local dynamics of microepidemics (Fraser et al., 2005; Hanage et al., 2006) and for longer time scales where recombination acts as a cohesive force keeping populations together(Fraser et al., 2007). However, there is limited knowledge about how varying levels of isolation in host organisms, such as human and different animal species (Fraser et al., 2009), might influence the evolutionary dynamics and lead to structured populations. Here we introduce a neutral model incorporating microepidemics and migration, which mimics a situation where ecological factors limit transmission between subpopulations. By comparing the model predictions with MLST data large scale genotyping surveys of four major human pathogens we find that for two species the population structure is well delineated by the neutral assumptions, while different types of deviations from the model predictions are observed for the remaining two.

The observed differences between *E. faecium* and *E. faecalis*, which colonize the gastrointestinal tract, are particularly interesting since mutation and recombination rates have been estimated to be similar for the two species based on both MLST and whole-genome data(de Been et al., 2013; Vos and Didelot, 2009). Moreover, they are responsible for roughly equal frequencies of nosocomial infections worldwide (Tedim et al., 2015; Willems et al., 2012). *E. faecalis* population structure bears the hallmarks of either a high rate of mutation or drift (or both). *E. faecalis* is known to colonize the vast majority of normal hosts within a population (Tedim et al., 2015), and therefore can be considered as part of the physiological commensal microbiota of humans and many other animals. Certainly, its population structure could be reflective of the evolutionary dynamics of a generalist organism which regularly experiences a high level of drift and gene flow between different host species.

On the basis of the predictions made by our model, *E. faecium* would need to have substantially higher recombination rate than *E. faecalis* to lead to the observed pattern of genotype relatedness under neutrality. Since there is evidence of the recombination rate not being substantially higher in *E. faecium*, the only possibility for the large genotype networks to arise under our neutral model would be unobserved population stratification. If unobserved sources experiencing very large clonal expansions contributed continuously to the hospital subpopulation of *E. faecium*, the expected allelic mismatch distribution would bear the characteristics of a subpopulation with high recombination rate (Supplementary Fig. 3, i). It is known that intensive farming and animal production practices provide opportunities for rapid clonal expansion of bacterial strains colonizing the animal hosts. Given the known connection between strains from domesticated animals and the hospital associated *E. faecium* (Lebreton et al., 2013; Willems et al., 2012), it is plausible that these clonal expansions could manifest themselves as connected networks in the human hospital subpopulation. However, the extensively connected network of *E. faecium* genotypes would still remain unlikely unless the rate of recombination was substantial. An alternative explanation for the extensive genotype relatedness is a marked deviation from neutrality, such that the connected strains represent either a subpopulation adapted to the hospital environment, consistent with previous studies(Lebreton et al., 2013; Willems et al., 2012), or an adaptation to different host subpopulations (Faith et al., 2015). Further dense sampling will be required to characterize mechanistically the role of hospital adaption for creating the observed relatedness patterns of *E. faecium* strains.

*S. aureus* and *S. epidermidis* frequently colonize the skin, soft tissue and the nares of human hosts, while also being ubiquitous in a range of animals. However, the overall population density and the proportion of human or animal hosts colonized by *S. epidermidis* largely exceed that of *S. aureus,* so that *S. epidermidis,* but not *S. aureus,* can be considered of a physiological commensal, part of the normal microbiota. The human *S. aureus* population is characterized by several genetically distinct clonal complexes, each sharing a single ancestral genotype. Such a population can arise under the neutral mutation/drift driven evolutionary trajectory combined with a high rate of localized transmission. In this scenario clonal complexes appear and proliferate for a time, to be replaced by others arising through genetic drift at the operational timescale of decades or longer. This has been previously described as an ‘epidemic clonal’ structure(Smith et al., 2000).

We may consider that *E. faecalis* and *S. epidermidis,* members of the normal microbiota, have an “endemic polyclonal structure”, where endemicity is assured by a highly frequent inter-host migration (both vertical and horizontal), resulting in a minimal adaptive stress in colonization of most hosts. On the contrary, *E. faecium* and *S. aureus* are less-adapted organisms to the generality of potential hosts, thus requiring local adaptation, and migration being dependent of this local success, an “epidemic clonal structure”. Obviously, in hospitals due to the homogenization of colonizable hosts (age, antibiotic exposure), and facilitation of host-to-host migration (hospital cross-colonization, microepidemics) *E. faecium* and *S. aureus* might appear as “locally endemic”, and therefore are expected to locally evolve towards a more complex population structure.

Both the commensal and hospital subpopulations of *S. epidermidis* display a pattern of genetic relatedness typical of a population where recombination is the dominant force generating population structure. An exception to this can be seen in the higher fraction of maximally distinct commensal genotypes, which could plausibly arise when novel strains infrequently migrate to the human commensal population from several non-overlapping zoonotic sources(Meric et al., 2015). However, our model was not able to accurately predict the persistence of the clonal complex structure observed for *S. aureus*, which may be reflecting a deviance from neutrality.

The complexities of within- and between-subpopulation strain dependence, and the extent of localized transmission and migration across ecological patch boundaries makes formal statistical inference about microepidemics and migration rates difficult. A particular challenge is that, when a population evolves within a drift dominated model, it is unlikely that reliable estimates of the parameters driving the population dynamics can be obtained, since observed outcomes of the population structure vary substantially. Similarly, as the consequences of migration events are dependent on other stochastically varying factors across the subpopulations, high migration rates may lead to a pattern of relatedness indistinguishable from those generated by low rates. It is possible that these issues could be resolved using coalescent-based models developed mainly for eukaryotic populations(Beerli and Felsenstein, 1999; Beerli and Felsenstein, 2001; Choi and Hey, 2011; Hey and Machado, 2003; Hey and Nielsen, 2004). However, robust generalization of such models is challenging due to the specific features of bacterial metapopulations which, in general, evolve by a complex combination of the stochastic forces of mutation, recombination, clonal expansion and host switches. Another obstacle for using coalescent-based methods is the large number of hosts that need to be explicitly considered in studies on large-scale bacterial pathogen populations.

It is evident that a limited number of neutrally evolving core genes, such as those typically used in the MLST typing schemes, limits the scope of models that can be fitted to genetic surveillance data. However, our results imply that some evolutionary scenarios would remain unidentifiable even if housekeeping loci were considered at the whole-genome scale, in particular if the data are mainly cross-sectional even if densely covering the host population. Hence, one of our main conclusions is that the optimal data for studying dynamics in this fashion are densely sampled longitudinal surveillance data covering evolutionary events at whole-genome level(Croucher et al., 2013). This highlights the importance of easy access online repositories of genomic variation as an extension of the currently existing MLST databases and that sample metadata should be an equally important focus of the data sharing principles. Using such a strategy in the near future may enable important model-based predictions about the dynamics of existing and emerging pathogens that pose a considerable global challenge for human and animal health.

## Acknowledgments

J.C., E.N., M.G. and M.S. were funded by the grant 251170 from Academy of Finland.

## Author contributions

J.C., E.N., M.G. developed and implemented the model, P.M. and M.S. provided additional expertise for the model development and analyses, J.C, G.M., S.K.S, T.C., F.B., W.V.S., R.W., E.F., W.P.H. provided data, biological expertise and interpretation, J.C., E.N., E.F. and W.P.H. wrote the manuscript. All authors approved the final manuscript.

